# The effects of the neonicotinoid imidacloprid on gene expression and DNA methylation in the buff-tailed bumblebee *Bombus terrestris*

**DOI:** 10.1101/590091

**Authors:** P.S.A Bebane, B.J. Hunt, M. Pegoraro, A.R.C Jones, H. Marshall, E. Rosato, E.B. Mallon

## Abstract

Neonicotinoids are effective insecticides used on many important arable and horticultural crops. They are nicotinic acetylcholine receptor agonists which disrupt the function of insect neurons and cause paralysis and death. In addition to direct mortality, there are numerous sublethal effects of low doses of neonicotinoids on bees. We hypothesize that some of these large array of effects could be a consequence of epigenetic changes in bees induced by neonicotinoids. We compared whole methylome (BS-seq) and RNA-seq libraries of the brains of buff tailed bumblebee *Bombus terrestris* workers exposed to field realistic doses of the neonicotinoid imidacloprid to libraries from control workers. We found numerous genes which show differential expression between neonicotinoid treated bees and control bees, but no differentially methylated cytosines in any context. We found CpG methylation to be focused mainly in exons and associated with highly expressed genes. We discuss the implications of our results for future legislation.

## Introduction

Neonicotinoids are effective insecticides used on many important arable and horticultural crops, most frequently as seed dressing. They are systemic, meaning they are absorbed by the plant and transported to all tissues where they remain active for many weeks or months. This protects all parts of the plant, but also means that neonicotinoids are found in the nectar and pollen of flowering crops such as oilseed rape, and hence are consumed by bees (Botías *et al.*, 2015). It has also emerged that they are commonly found contaminating nectar and pollen of wild flowers growing on arable farmland, providing additional exposure of bees and other pollinators (Botías *et al.*, 2015; David *et al.*, 2016).

Neonicotinoids are nicotinic acetylcholine receptor agonists which disrupt the function of insect neurons and cause paralysis and death. In addition to direct mortality, laboratory and field studies have documented numerous sublethal effects of low doses of neonicotinoids on both honeybees and bumblebees (e.g. Whitehorn *et al.* 2012; Rundlöf *et al.* 2015, reviewed in Pisa *et al.* 2015). Sublethal effects at the individual level include reduced fecundity of queens, reduced fertility in males, impaired immune response, impaired navigation and learning, reduced pollen collection and reduced food consumption. Collectively, these effects result in reduced colony growth and colony reproduction performance. The breadth of the effects of neonicotinoids on bees suggests that neonicotinoids have multiple modes of action beyond their designed direct impact on neurotransmission, for example their impact on immune signalling (Prisco *et al.*, 2013).

We hypothesize that some of these effects could be a consequence of epigenetic changes induced by neonicotinoids. Epigenetics is defined as the stable and heritable change in gene expression without any change in the DNA sequence (Goldberg *et al.*, 2007). Environmental contaminants have been found to affect the epigenetics of a diverse range of animal species from water fleas to polar bears (Head, 2014) and include metals, endocrine disrupting compounds, air pollution, persistant organic pollutants and pesticides (Vandegehuchte and Janssen, 2014), but much ecotoxicology research is centred on a direct link between exposure and response (Head, 2014). Epigenetic changes have the potential to weaken that link, with effects possibly manifesting much later in life or in subsequent generations. Thus if pesticide-induced epigenetic changes were shown to be heritable in bees this would have implications for future ecological risk assessment.

In social insect research the role of DNA methylation, an epigenetic marker primarily involving the addition of a methyl group to a cytosine, has come under increasing scrutiny in recent years (Foret *et al.*, 2009; Lyko *et al.*, 2010; Glastad *et al.*, 2013; Amarasinghe *et al.*, 2014; Glastad *et al.*, 2016; Patalano *et al.*, 2015; Libbrecht *et al.*, 2016; Standage *et al.*, 2016; Rehan *et al.*, 2016; Glastad *et al.*, 2017; Arsenault *et al.*, 2018). Methylation has also been implicated in important effects on the biology of bees, including the control of reproductive status (Kucharski *et al.*, 2008; Amarasinghe *et al.*, 2014) and memory (Biergans *et al.*, 2012), behaviours shown to be affected by neonicotinoids (Williams *et al.*, 2015; Stanley *et al.*, 2015), although in the case of reproduction the link between methylation and social insect reproduction is controversial (Herb *et al.*, 2018; Patalano *et al.*, 2015; Libbrecht *et al.*, 2016). DNA methylation has been linked with alternative splicing in a number of insect species (Lyko *et al.*, 2010; Li-Byarlay *et al.*, 2013; Glastad *et al.*, 2016; Arsenault *et al.*, 2018), and with histone modifications in the ant *Camponotus floridanus* (Glastad *et al.*, 2015). In mammals, methylation on gene promoters leads to a reduction in gene expression. The effect of methylation on gene expression in insects is less well understood (Pegoraro *et al.*, 2017), though high levels of methylation have been associated with highly and stably expressed genes (Foret *et al.*, 2012; Bonasio *et al.*, 2012; Wang *et al.*, 2013), while in honeybees hypomethylated genes are associated with caste-specific expression (Elango *et al.*, 2009; Libbrecht *et al.*, 2016; Marshall *et al.*, 2019). Gene expression differences due to neonicotinoid exposure have been found in honeyebee larval workers, adult workers and queens (Derecka *et al.*, 2013; Aufauvre *et al.*, 2014; Christen *et al.*, 2016; Chaimanee *et al.*, 2016; Christen *et al.*, 2018).

In this study we use whole genome bisulfite sequencing (WGBS/BS-seq) and RNA-seq on brain tissue of neonicotinoid exposed and control *Bombus terrestris* workers in order to elucidate the effects of the neonicotinoid imidacloprid on the gene expression and methylation status of bumblebee workers.

## Materials and Methods

### Beekeeping, experimental design and brain dissection

Six colonies of *Bombus terrestris audax* were purchased from Agralan, UK. Each colony contained a queen and on average ten workers and a small amount of brood. They were kept in wooden nest boxes and maintained under red light at 26°C and 60% humidity on a diet of 50% v/v glucose/fructose apiary solution (Meliose-Roquette, France) and pollen (Percie du set, France) (Amarasinghe *et al.*, 2014). Three colonies were used for the RNA-seq experiment and the other three for the BS-seq experiment (Figure S1).

Groups of 5 callow workers born on the same day were reared in Perspex boxes (18.5 cm x 12.5cm x 6.5cm). Boxes were then randomly assign to control or treated groups. The control group was fed *ad libitum* with 50% v/v apiary solution for six days whereas the treated group was fed *ad libitum* with a 10ppb imidacloprid (SIGMA-ALDRICH) 50% v/v apiary solution, a field-realistic sub-lethal dose (Cresswell, 2011; Blacquière *et al.*, 2012). After a six day chronic exposure period (Cresswell, 2011) the bees were anesthetized on ice at 4°C. The brains were dissected in phosphate buffered saline (PBS) and immediately frozen in liquid nitrogen and stored at −80°C. Their ovaries were checked for development to ensure that only non-reproductive workers were used (Amarasinghe *et al.*, 2014; Harrison *et al.*, 2015).

### BS-seq

#### Genomic DNA extraction, sequencing and mapping

Six libraries were prepared (3 colonies, control and treatment). For each colony, 10 boxes were reared (5 control and 5 treatment). Each library was generated from 12 pooled brains of non-reproductive workers taken at random from the relevant boxes for a total of 72 brains. Genomic DNA was extracted, using QIAGEN QIAamp DNA Micro Kit following the manufacturer’s instructions. The concentration of genomic DNA was measured using a Qubit® dsDNA BR Assay Kit (ThermoFisher Scientific, USA) and Nanodrop. Sequencing was performed on a HiSeq 2000 machine (Illumina, Inc.) at the Beijing Genomics Institute (BGI), generating 100-bp paired-end reads.

Poor quality reads were removed using fastQC v0.11.2 (Andrews, 2010) and adapters trimmed using cutadapt V1.11 (Martin, 2011) and trimmomatic V0.36 (Bolger *et al.*, 2014). Bismark v0.18.1 (Krueger and Andrews, 2011) was used to align the reads to the Bter_1.0 genome (Refseq accession no. GCF_000214255.1 (Sadd *et al.*, 2015)), remove PCR artifacts and extract methylation calls in CpG, CHH and CHG contexts (where H represents adenine, thymine or cytosine). The cytosine report files from Bismark and the *B. terrestris* annotation file (GCF_000214255.1) were combined using the sqldf library (Grothendieck, 2017) in R v3.4.0 (R Core Team, 2014) to generate the distribution of methylated Cs over genomic features. Cytosines with less than 10X coverage were excluded. For each cytosine the proportion of methylation reads over total reads was calculated.

#### Methylation differences between treatments

Differential methylation analysis was performed using methylKit (Akalin *et al.*, 2012). Bismark cytosine reports were filtered to exclude loci with extreme low or high coverage (< 10 or > 500 reads) and those not covered in all samples. A mixture of binomial model (Cheng and Zhu, 2014) was used to make per-loci methylation status calls and only loci identified as methylated in at least one sample were tested. A logistic regression test was applied using overdispersion correction, controlling for colony as a covariate, and adjusting p-values for multiple testing using the SLIM method. A minimum change in methylation between treatments of 10% was used to filter results.

### RNA-seq

#### RNA extraction and Illumina sequencing

Eighteen libraries were prepared (three colonies, three replicates per colony, two conditions). For each colony, 6 boxes were reared (3 control and 3 treatment). Each library was generated from 3 pooled brains of non-reproductive workers taken from the relevant boxes, for a total of 54 brains. Total RNA was isolated utilizing the GenElute Mammalian Total RNA Miniprep Kit. DNA and RNAase activity was eliminated using (Sigma-Aldrich DNase I treatment kit) following the manufacturer’s instruction. RNA concentration and integrity were determined by Bioanalyzer using the RNA Nano Kit (Agilent Technologies). From each sample we isolated an average of 0.8 mg of RNA. Two samples appeared degraded and were not used. Nine control and seven treated samples were prepared and sequenced on HiSeq 200 (Illumina, Inc.) at Beijing Genomics Institute (BGI) and 100-bp paired-end reads were generated.

BGI removed adaptor sequences, contamination and low-quality reads from raw data. Base calling and quality scoring of the raw reads were visualized using fastQC v 0.11.2 (Andrews, 2010). The clean reads for each sample were aligned to the reference genome Bter_1.0 genome (Refseq accession no. GCF_000214255.1 (Sadd *et al.*, 2015)) using Hisat2 v2.0.4 (Kim *et al.*, 2015) with default parameters. The output sam file was sorted and converted to a bam file using samtools (Li *et al.*, 2009). Aligned reads were assembled and quantified using the assembler stringtie v1.3.3b (Pertea *et al.*, 2015).

#### Differential gene expression analysis

A table of raw counts was generated using a Python script (https://github.com/gpertea/stringtie/blob/master/prepDE) and analysed using DESeq2 (Love *et al.*, 2014) in R v3.4.0 (R Core Team, 2014) to estimate differentially expressed genes using an FDR-adjusted p-value threshold of 0.05 and controlling for colony effects. Genes with less than 10 reads were discarded from analysis. The normalized read counts were log_2_ transformed. The quality of replicates was assessed by plotting read counts of samples against one another and assessing the dispersion and presence of any artefacts between samples (Rich *et al.*, 2018). A principal-component analysis was performed to visualize diversity between samples within treatment and between condition.

### GO term enrichment and KEGG analysis

A list of GO terms for the bumblebee were made by annotating the transcriptome using trinotate (default settings) (Hébert *et al.*, 2016) and blast2GO (against RefSeq) (Conesa *et al.*, 2005). These lists were combined, using the pipeline implemented in Amar *et al.* 2014 with a K value of 1. A hypergeometric test was applied and significant GO terms identified after BH correction (p corrected < 0.05) (Benjamini and Hochberg, 1995) using GOstats (Falcon and Gentleman, 2007), with all RNA features in the bumblebee genome used as a background (GCF_000214255.1). We filtered these to only those terms present in three or more DEGs and used REVIGO (Supek *et al.*, 2011) to cluster and visualise enriched GO terms, selecting the whole UniProt database and SimRel semantic similarity measure.

The clusterprofiler R package (version 3.8.1) (Yu *et al.*, 2012) identified differentially expressed genes associated with KEGG pathways using the whole UniProt database. A hypergeometric test was applied and significant KEGG pathways were identified after BH correction (qvalue < 0.05) (Benjamini and Hochberg, 1995).

## Results

### Methylation analysis

The overall sequence alignment rate was 67.21% 1.53% (mean standard deviation). The proportion of methylated cytosine reads calculated by Bismark were 0.53% 0.05% for CpGs, 0.37% 0.05% for CHGs, 0.38% 0.07% for CHHs and 0.4% 0.06% for CNs or CHNs ((H = A, C, or T). While insect methylation levels are often low (Glastad *et al.*, 2017) these methylation levels are lower even than in the honey bee, *Apis mellifera*, estimated at ~1% at the genome level using similar metrics (Feng *et al.*, 2010; Bewick *et al.*, 2017). In a CpG context, across all samples, 0.15% 0.03 % of loci with a minimum coverage of 10 reads were considered methylated by the mixture of binomial model. The distribution of CpG methylation shows a mild bimodal distribution with the vast majority of sites being not or only modestly methylated and a few fully methylated (Figure S2 A). Methylated CpGs are more abundant in coding regions (seven fold) and exons (five fold) than introns (Figure 1 A). Non-CpG per-loci methylation levels were reported as less than 0.001% by the mixture of binomial model. This, in conjunction with the uniformity of non-CpG methylation across genomic features (Figure 1 B,C), led to the conclusion that such levels were indistinguishable from error and as such were excluded from subsequent analysis.

**Figure 1:**
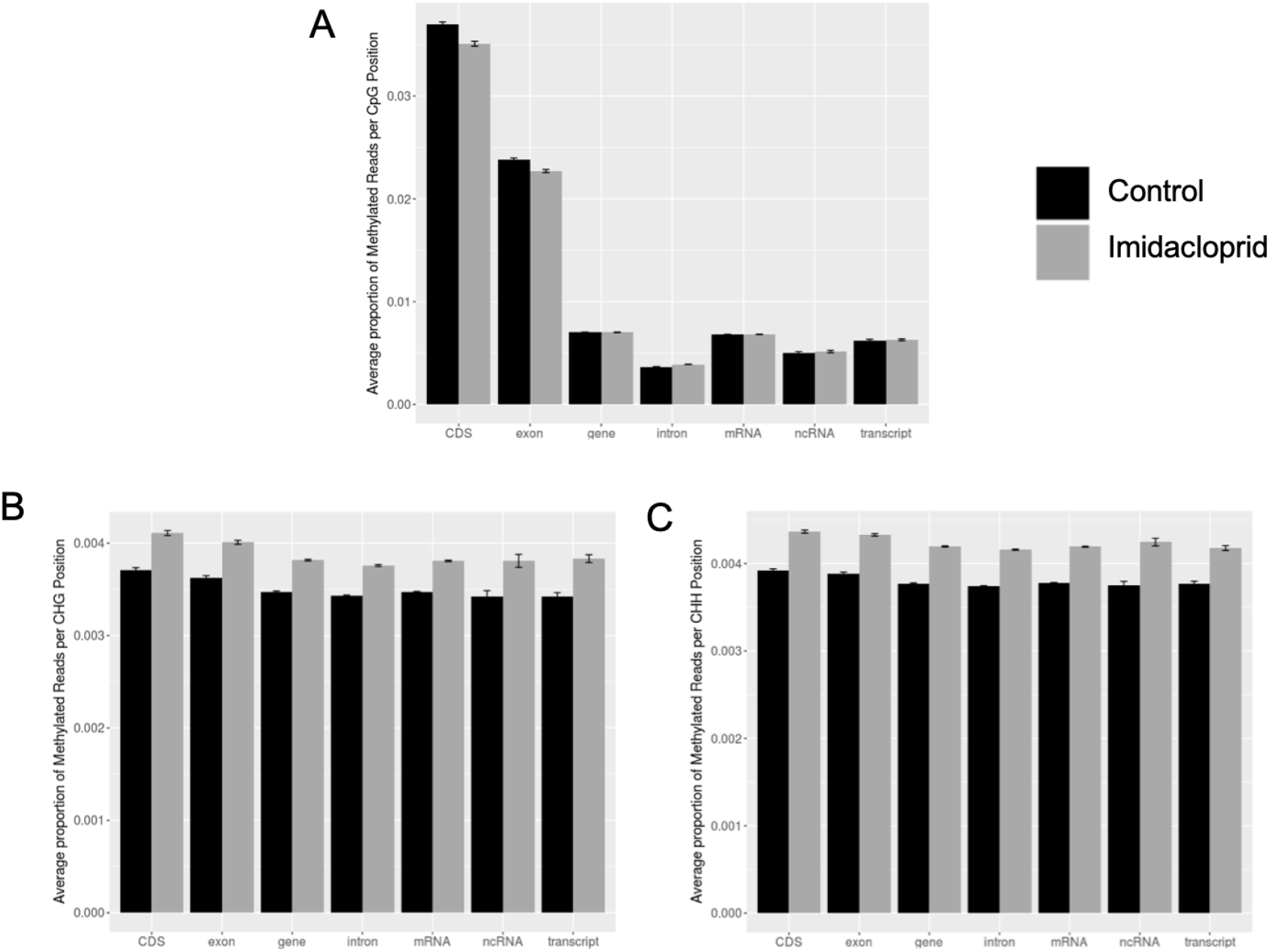
Methylated Cs distribution. Average proportion of methylation reads SD per CpG (**A**), CHG (**B**) and CHH (**C**) positions over genomic features. Control samples in black and Neo treated samples in grey.

#### Methylation differences between control and neonicotinoid treated samples

In total 4,424,986 loci were analysed using the mixture of binomial model, which subsequently identified 6,080 sites to test. No differentially methylated loci were identified using logistic regression at a q-value of 0.05 or 0.1. MethylKit includes an option to pool replicates into single control/treatment samples and use Fisher’s exact test; using this approach we identified a small number of differentially methylated CpGs at q-value < 0.1, including loci within *histone-lysine N-methyltransferase 2C*, *histone acetyltransferase p300*, *CXXC1* (a transcriptional activator that binds to unmethylated CpGs), and genes involved with axon formation (supplementary data, diff_meth_fisher).

### Expression analysis

Alignment rate to the genome was 93.6% (92.1 to 94.1) and after filtering a total of 10,772 genes were analysed. All libraries from the same treatment showed low variation in their gene expression patterns (Figure S3, S4).

#### Differential expression

A total of 405 genes were differentially expressed: 192 genes upregulated and 213 downregulated in neonicotinoid samples compared to controls (see supplementary data: differentially_expressed_genes).

Four cytochrome P450 (CYP) genes were differentially expressed, two upregulated and two downregulated. Upregulated genes in neonicotinoid treated bees also include *apyrase* that hydrolyzes ATP to AMP, the neuropeptide receptor *pyrokinin-1 receptor* and *ionotropic receptor 25a* that is involved in circadian clock resetting in *Drosophila* (Chen *et al.*, 2015). Downregulated genes include *neurexin*, involved in synaptic formation and maintenance, *peptide methionine sulfoxide reductase*, involved in repair of oxidation-damaged proteins, and a number of genes related to photoreceptor function. Three genes belonging to the homeotic box gene (Hox) family were downregulated in neonicotinoid treated bees. *lethal(2)essential for life (Efl21)* displayed the highest down regulation. We found 105 enriched biological process GO terms (BH corrected p < 0.05) associated with differential gene expression (supplementary data: expression_GO), subsequently clustered using REVIGO to 58 terms (Figure S5). Many of the most significantly enriched terms were associated with energy reserve metabolism. Also enriched were terms associated with apoptotic processes, apoptotic cell clearance, immune effector processes, cell death and response to chemical stimulus. No KEGG pathways were over represented for differentially expressed genes (q < 0.05).

### DNA methylation - Expression correlation

We calculated the average percentage of methylated reads per gene for the most differentially expressed genes (log_2_ fold-change > 0.5 or < −0.5) and non-differentially expressed genes (Figure 2), fitting a generalized linear model (GLM) with a quasi binomial error distribution with treatment (control vs neonicotinoid) and expression state (DEG vs. non-DEG) as independent variables. There was no significant interactions between the independent variables (interaction model versus main effects only model: *χ*^2^ = −0.014, d.f. = 1, p = 0.82). For CpGs, non-differentially expressed genes had more methylation than differentially expressed genes (z_1,19673_=4.641, p<0.001). There was no significant treatment effect on methylation levels (z_1,19673_=−0.772, p=0.692).

**Figure 2:**
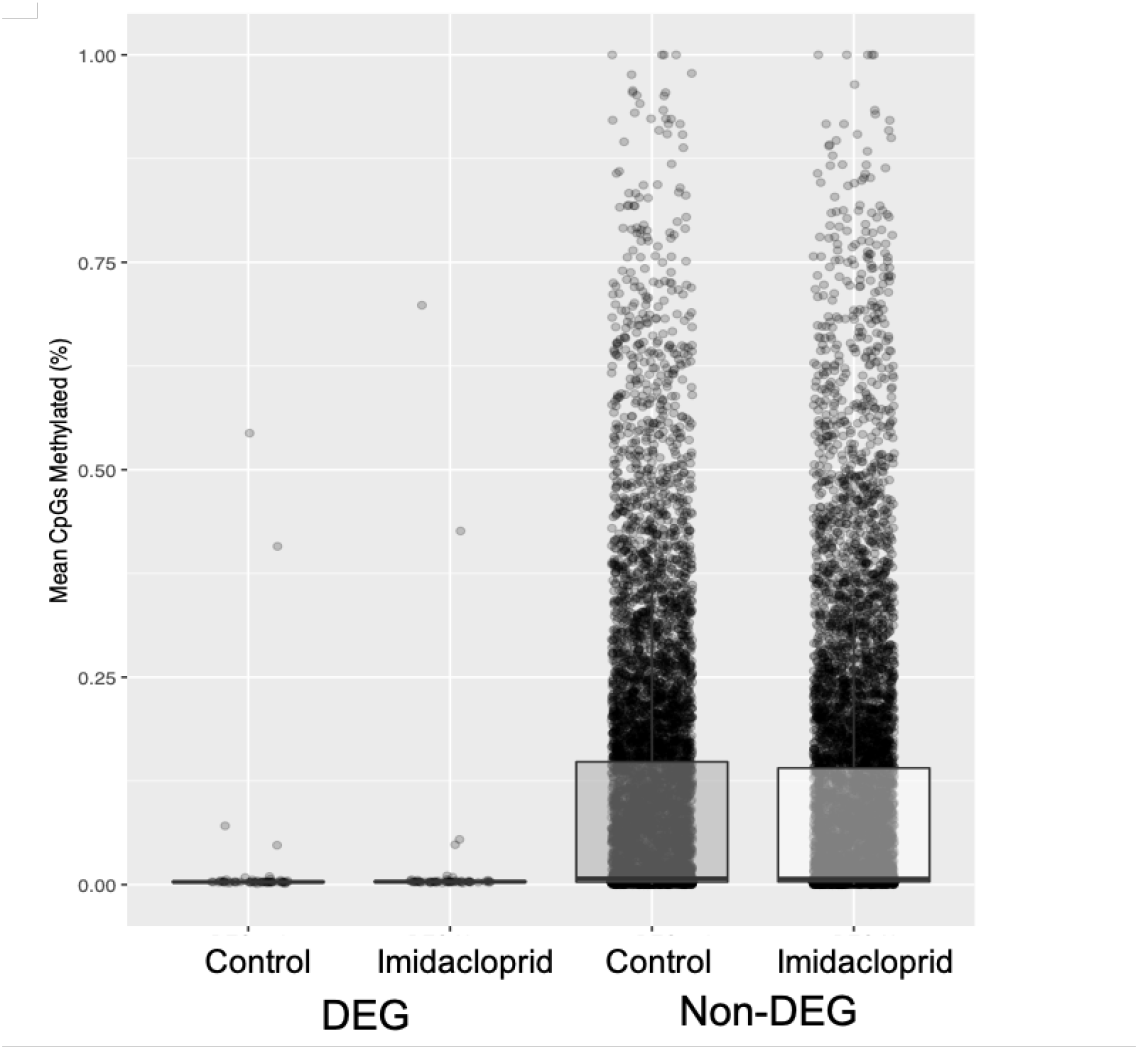
Average percentage of methylated CpG per gene. Differentially expressed genes (DEG) and non differentially expressed genes (nonDEG) are plotted separately. Dots represent genes.

To have a more fine scale understanding of the correlation between methylation and expression, we plotted mean proportion of methylation per gene against ranked expression level (log_10_fpkm per gene) in 100 bins (from low to high) (Figure 3) fitting a linear model with treatment and expression level as independent variables. There was no significant interaction between expression’s and treatment’s effects on methylation (interaction model versus main effects only model: F_1,189_ = 1.0347, p = 0.3104). We found a significant association between expression and methylation (F_1,189_ = 281.654, p = < 2 x 10^−16^). Neonicotinoid treated bees had comparable levels of CpG methylation to control bees (F_1,189_ = 1.8125, p = 0.1798).

**Figure 3:**
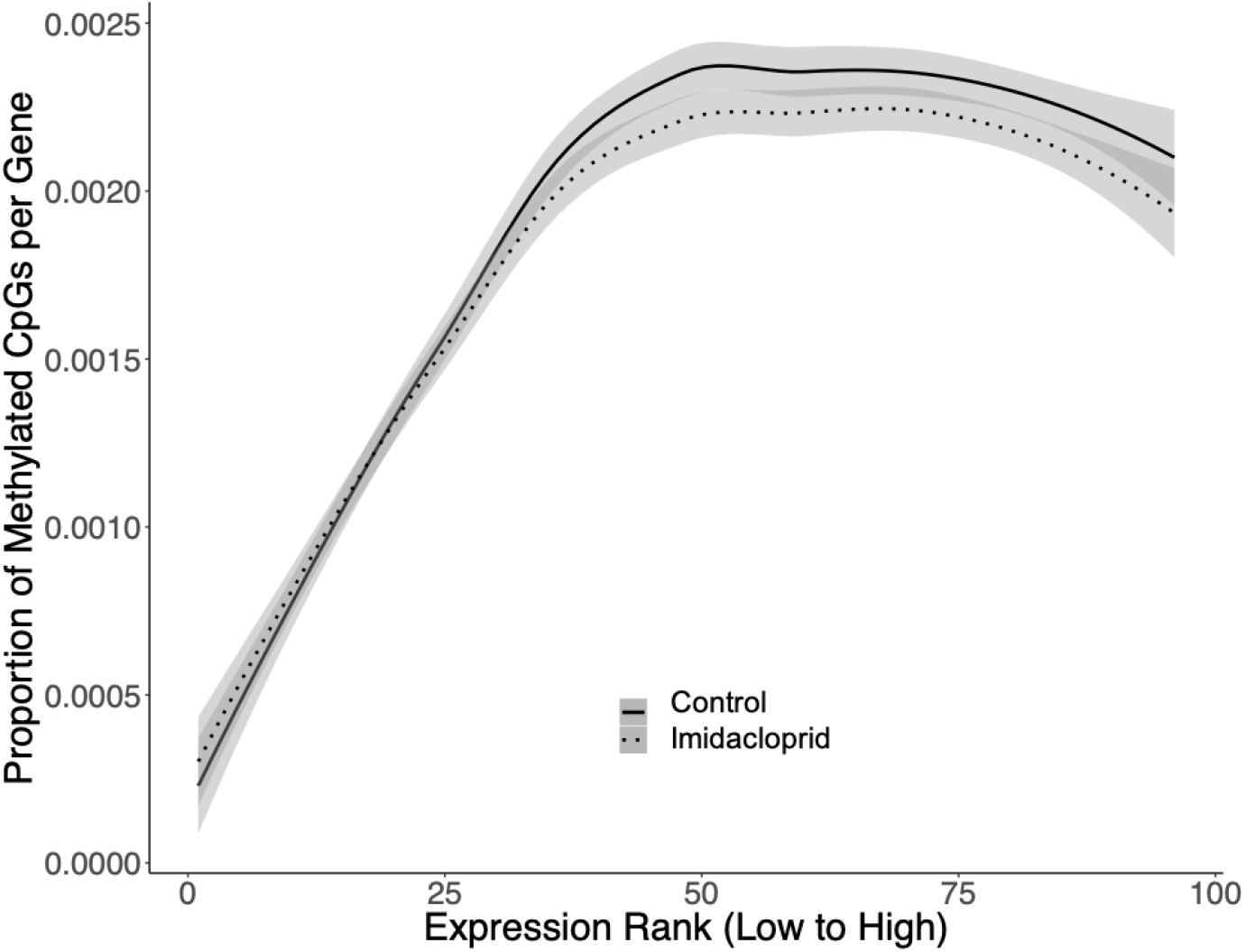
The proportion of methylated CpGs is plotted against gene expression rank. One hundred “bins” of progressively increasing level of expression were generated and genes with similar level of expression have been grouped in the same bin. Solid lines represent control samples and dotted lines neonicotinoid treated samples. The grey shading represents 95% confidence intervals.

## Discussion

We found numerous genes which show differential expression between bees treated with field realistic doses of the neonicotinoid imidacloprid and control bees. We found CpG methylation to be focused in exons, and high CpG methylation was associated with highly expressed genes, but no differentially methylated loci were detected between treatments. Non-differentially expressed genes had higher methylation levels than differentially expressed genes.

Four cytochrome P450 (CYP) genes were identified as differentially expressed, in line with other studies assessing the impact of insecticides on honeybees (Shi *et al.*, 2017; Li *et al.*, 2017; Derecka *et al.*, 2013; Wu *et al.*, 2017; Christen *et al.*, 2018). Two were upregulated (CYP6k1 and 4c3) and two downregulated (28d1 and 9e2). CYP6, 9 and 28 genes are linked to xenobiotic metabolism and resistance to insecticides (Feyereisen, 2006) and CYP6 genes specifically have been found to be upregulated in honeybees after treatment with sublethal doses of the neonicotinoid Thiamethoxam (Shi *et al.*, 2017), as has CYP4C1 after treatment with the neonicotinoid Clothianidin (Christen *et al.*, 2018). The CYP9Q subfamily were recently shown to be responsible for bee sensitivity to neonicotinoids (Manjon *et al.*, 2018).

The identification of differentially expressed genes associated with synaptic transmission (supplementary data: expression_GO) is to be expected, given that we used brain tissue and given the known target effects of neonicotinoids. The identification of a downregulated *neurexin* gene aligns with the results of Shi *et al.* (2017). The effect seen here on metabolic pathways has also been found in honeybees, with GO term enrichment for catabolic carbohydrate and lipid metabolism (Christen *et al.*, 2018). These authors suggested that due to the intensive energy demands of the brain, negative effects on metabolic pathways could affect brain function and therefore behaviour. During the review period a further study was published examining gene expression changes in *B. terrestris* after exposure to neonicotinoids, again showing changes in carbohydrate and lipid metabolism (Colgan *et al.*, 2019). *Efl21*, the most downregulated gene identified, has been found to be involved in foraging behaviour in bees (Hernández *et al.*, 2012), a potential genetic link to the findings of Mommaerts *et al.* (2009). Impaired foraging has implications for pollination, reproduction and overall colony survival. Downregulation of carbohydrate metabolism pathways has also been shown in honeybee larvae (Derecka *et al.*, 2013; Wu *et al.*, 2017). Also downregulated were three hox genes. This may be indicative of an impaired immune system, as hox genes have been found to play a role in invertebrate innate immune responses (Uvell and Engström, 2007; Irazoqui *et al.*, 2008). Hox genes have been found to be downregulated in response to insecticide treatment in honeybees (Aufauvre *et al.*, 2014). The bumblebee visual system may also be impacted by imidacloprid treatment, given the downregulation of genes such as protein scarlet, protein glass and ninaC.

No differentially methylated loci between control and treatment were identified using a logistic regression model, and we suggest that if acute neonicotinoid exposure does alter methylation status in *B. terrestris* it is subtle and the data reported here may be underpowered to detect it due to low per-sample coverage. A small number of differentially methylated loci were identified by pooling replicates and using Fisher’s exact test (supplementary data: diff_meth_fisher), but unlike logistic regression this approach cannot control for covariates and the results should be treated with caution. Using this approach a CpG loci in *CXXC-type zinc finger protein 1* was identified as hypermethylated in neonicotinoid-treated bees; this gene also was upregulated in that group. In mammals, CXXC1 is a transcriptional activator that binds to unmethylated CpGs to regulate gene expression (Shin Voo *et al.*, 2000). Other loci identified by pooling were located within *histone acetyltransferase p300* and *histone-lysine N-methyltransferase 2C*. These findings raise the possibility that neonicotinoids may have a more detectable effect over a longer period through a cascade of epigenetic processes. A study on the effects of imidacloprid on bumblebees found no effect on mortality or reproduction over 11 weeks using 10 ppb when workers were not required to forage for food, while 20 ppb affected mortality and foraging was impaired at both doses (Mommaerts *et al.*, 2009). It may therefore be that a higher dose or longer exposure time might have a detectable impact on CpG methylation, and further work investigating chronic rather than acute exposure to imidacloprid at different doses would be valuable. Also worthy of investigation is the potential effect on epigenetic processes other than DNA methylation, such as histone modification, which has been found to have a similar, but non-redundant, association with gene expression in the ant *Camponotus floridanus* (Glastad *et al.*, 2015).

We found patterns of CpG methylation to be in line with other insect species. It is mainly focused in exons (Glastad *et al.*, 2017), and high CpG methylation was associated with highly expressed genes (Figure 3) (Arsenault *et al.*, 2018; Bonasio *et al.*, 2012; Glastad *et al.*, 2013; Libbrecht *et al.*, 2016; Patalano *et al.*, 2015; Wang *et al.*, 2013), and non-differentially expressed genes showed higher levels of methylation (Glastad *et al.*, 2013, 2016; Libbrecht *et al.*, 2016; Sarda *et al.*, 2012). As well as inducing no changes in methylation at individual loci, neonicotinoids appear to have no effect on overall levels of CpG methylation (see Figures 2 and 3). This failure to identify methylation differences between experimental groups is consistent with findings of robust methylation between castes in various insects (Hunt *et al.*, 2010) but contrasts with studies finding differences resulting from removal of maternal care (Arsenault *et al.*, 2018), or within castes with differing reproductive status (Marshall *et al.*, 2019).

Non-CpG methylation plays a role in gene silencing in flowering plants (Stroud *et al.*, 2014) and to a lesser extent, in mammals (Dyachenko *et al.*, 2010). In this study, while we identified a very small number of loci showing methylation in CHG/CHH contexts we could not exclude the possibility that much of it was noise, as bisulfite sequencing is prone to false positives from sources such as incomplete bisulfite conversion, miscalled bases and SNPs. Overall, we conclude that there is no notable methylation of non-CpG cytosines in *B. terrestris*, as with the honeybee (Lyko *et al.*, 2010) and *Nasonia vitripennis* (Wang *et al.*, 2013). In contrast to the preponderance of CpG methylation in exons, we found that CHH and CHG methylation was uniformly spread throughout genes (Figure 1) a pattern which would be consistent with the idea that there is no significant methylation in these contexts.

Recently, it has become clear that epigenetics can play a role in the interplay between man-made chemicals and natural ecosystems, and their constituent species (Vandegehuchte and Janssen, 2014). Hymenopteran insects (ants, bees and wasps) are ideal models to study this. They are both strongly affected by man-made chemicals and are important emerging models for epigenetics, with a number of species with relatively small genomes showing a confirmed role for methylation in their biology (Glastad *et al.*, 2011; Weiner and Toth, 2012; Welch and Lister, 2014; Yan *et al.*, 2014).

However, on the evidence of this study, imidacloprid does not appear to have epigenetic effects, at least through DNA methylation. This finding is important in the context of future legislation for pesticide control, as it is evidence suggesting a potential lack of transgenerational effects on *B. terrestris* with the use of imidacloprid.

## Acknowledgements

Thanks to Dr. Swidbert Ott and Prof. Dave Goulson for discussions. EBM, BJH and MP were funded by NERC grant NE/N010019/1. PSAB was supported by a scholarship from the Human Capacity Development Program (Koya University - Iraq). HM was supported by a NERC CENTA DTP studentship. ARCJ was supported by a BBSRC MIBTP DTP studentship. This research used the ALICE High Performance Computing Facility at the University of Leicester.

## Data accessibility

All sequencing data related to this project can be found under NCBI BioProject PRJNA524132.

## Authors’ contributions

EBM, ER and PSAB designed the study. PSAB carried out the experiments. PSAB, BJH, MP, ARCJ and HM analysed the data. MP, PSAB and EBM wrote the initial draft. All authors were involved in redrafting.

## Supplementary material

Supplementary figures are availible in the supplementary figures file. Supplementary data is available at https://doi.org/10.6084/m9.figshare.6796802.

## Notes

https://figshare.com/articles/supplementary_data/6796802/6

