## Supplementary figures for "The effects of the neonicotinoid imidacloprid on gene expression and DNA methylation in the buff-tailed bumblebee *Bombus terrestris*"

1) Department of Genetics and Genome Biology
University of Leicester
Leicester

2) School of Natural Sciences and Psychology
John Moores University
Liverpool

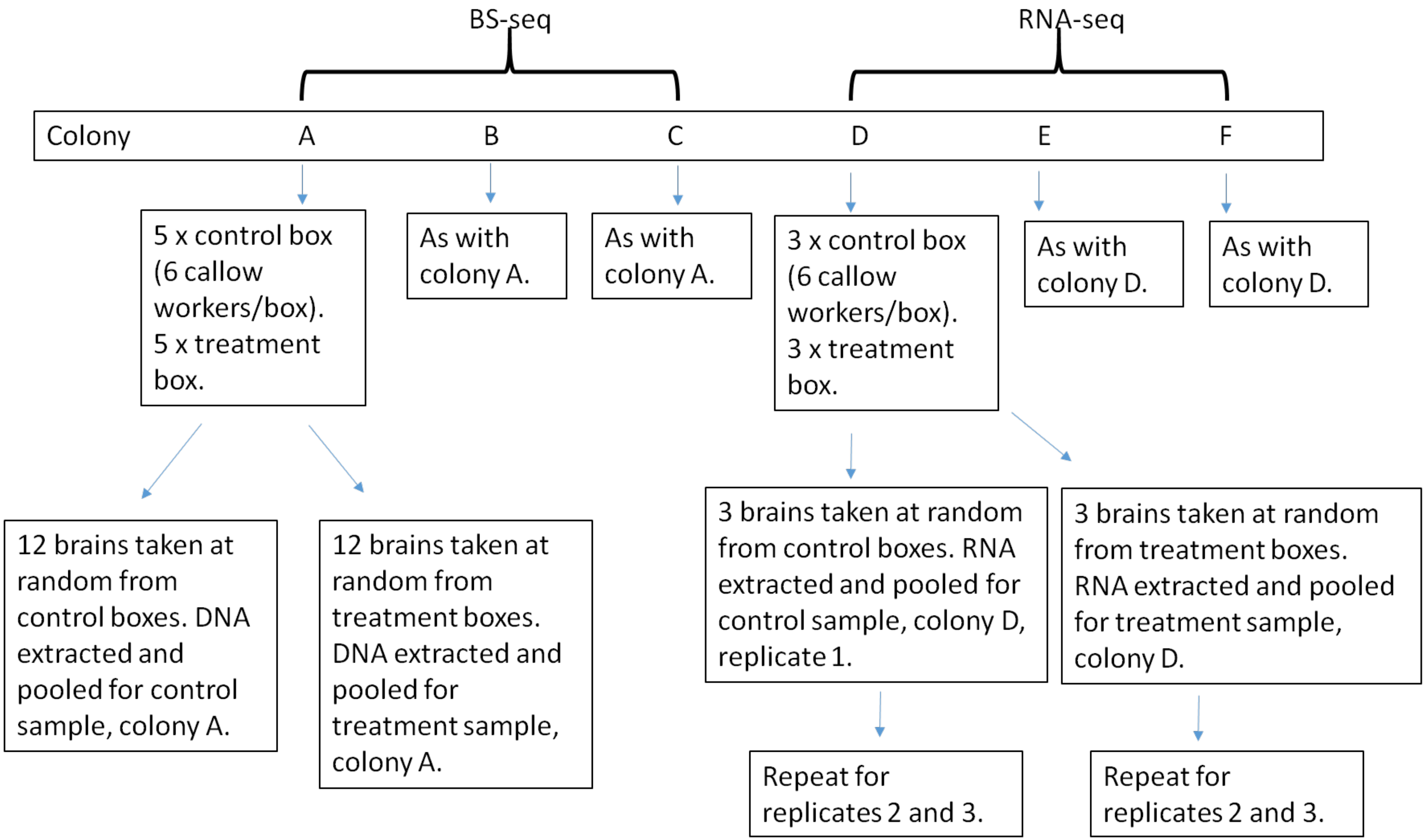


Figure S1: Sample collection for BS-seq and RNA-seq projects.

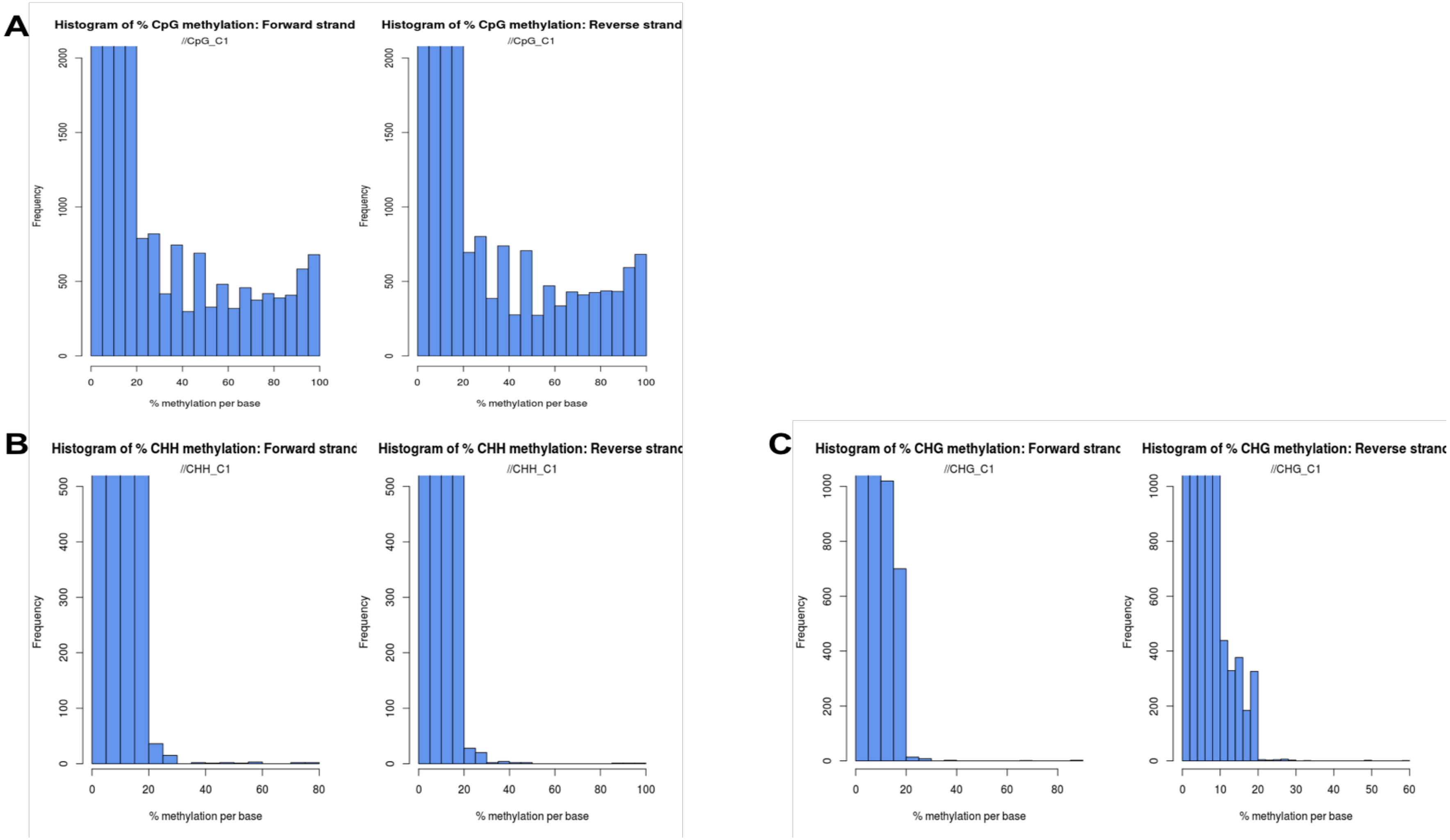


Figure S2: Examples of histogram of percentage methylation per cytosine. Histogram of % methylation per CpG (A), CHH (B) and CHG (C) for a control sample (C1). CpG % methylation shows a mild bimodal distribution with most sites modestly methylated and few fully methylated. Contrary to CpG, most CHH and CHG are modestly methylated. Both forward and reverse reads are reported.

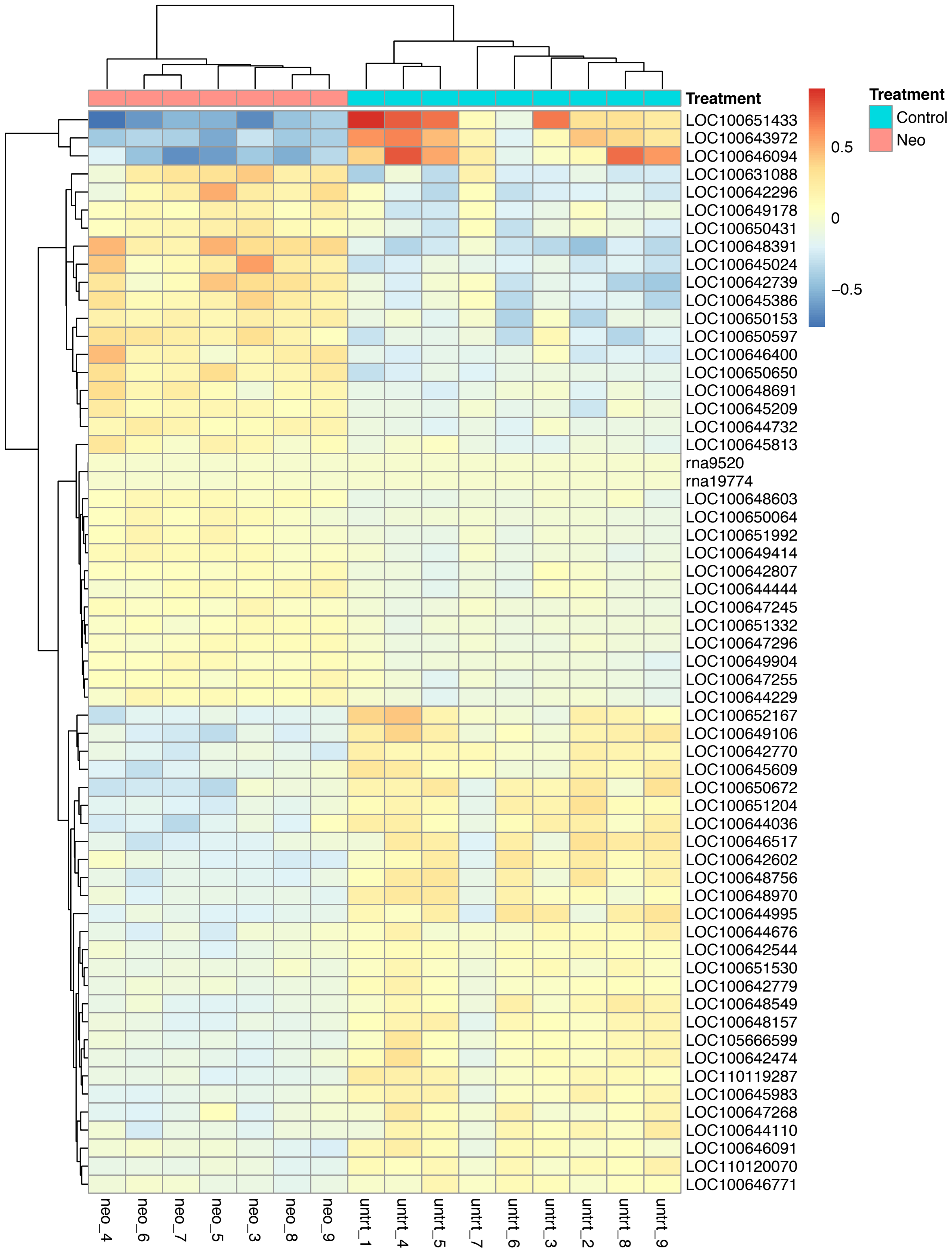


Figure S3: Heatmap tree showing 30 genes. 15 genes that are up regulated and 15genes are down regulated. The yellow means high level of gene expression while blue shows reduced relative gene expression

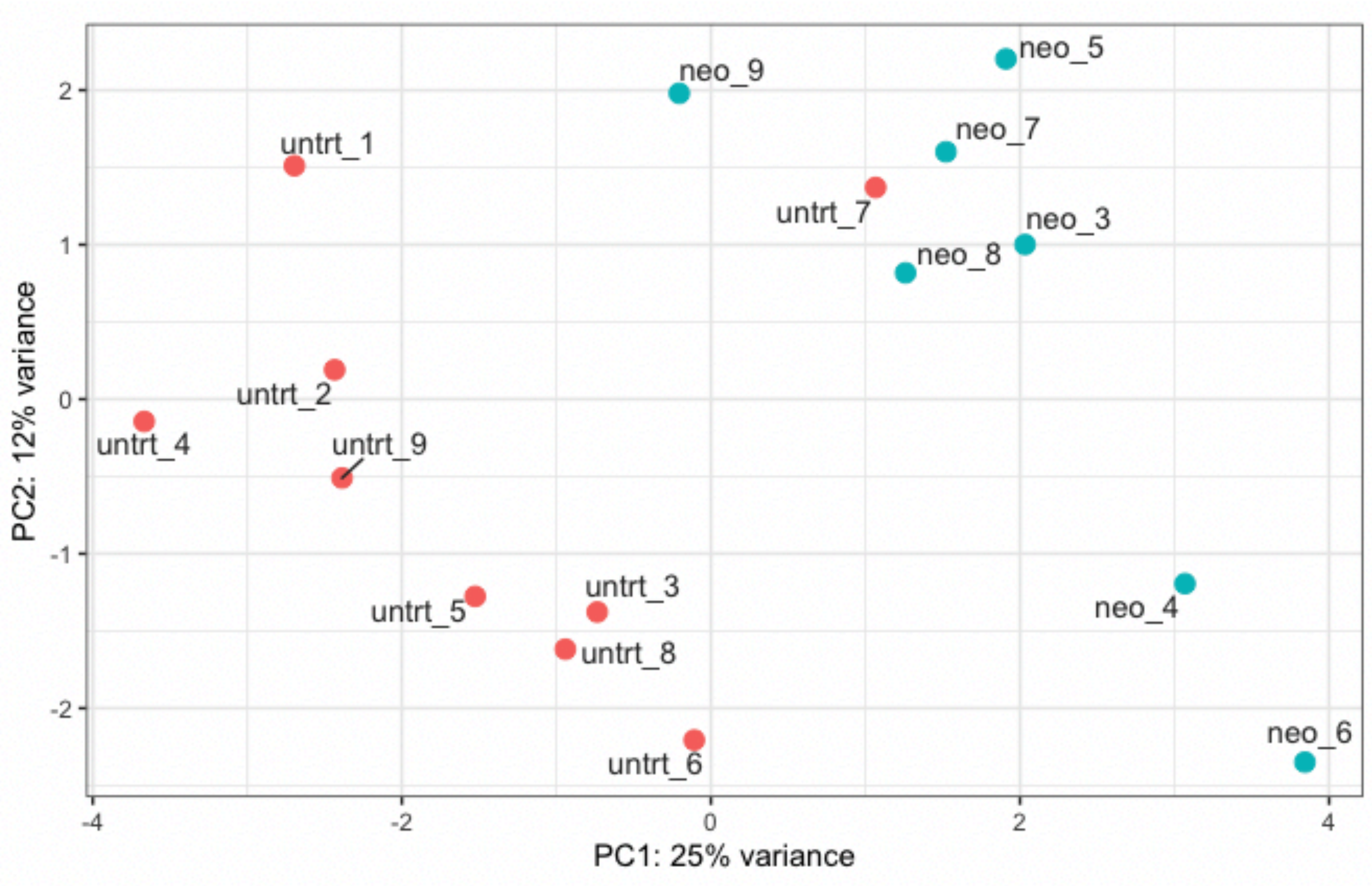


Figure S4: PCA plots showing correlation between samples and distance between imidacloprid and control condition. Red dots representing Neo samples and blue points representing control samples

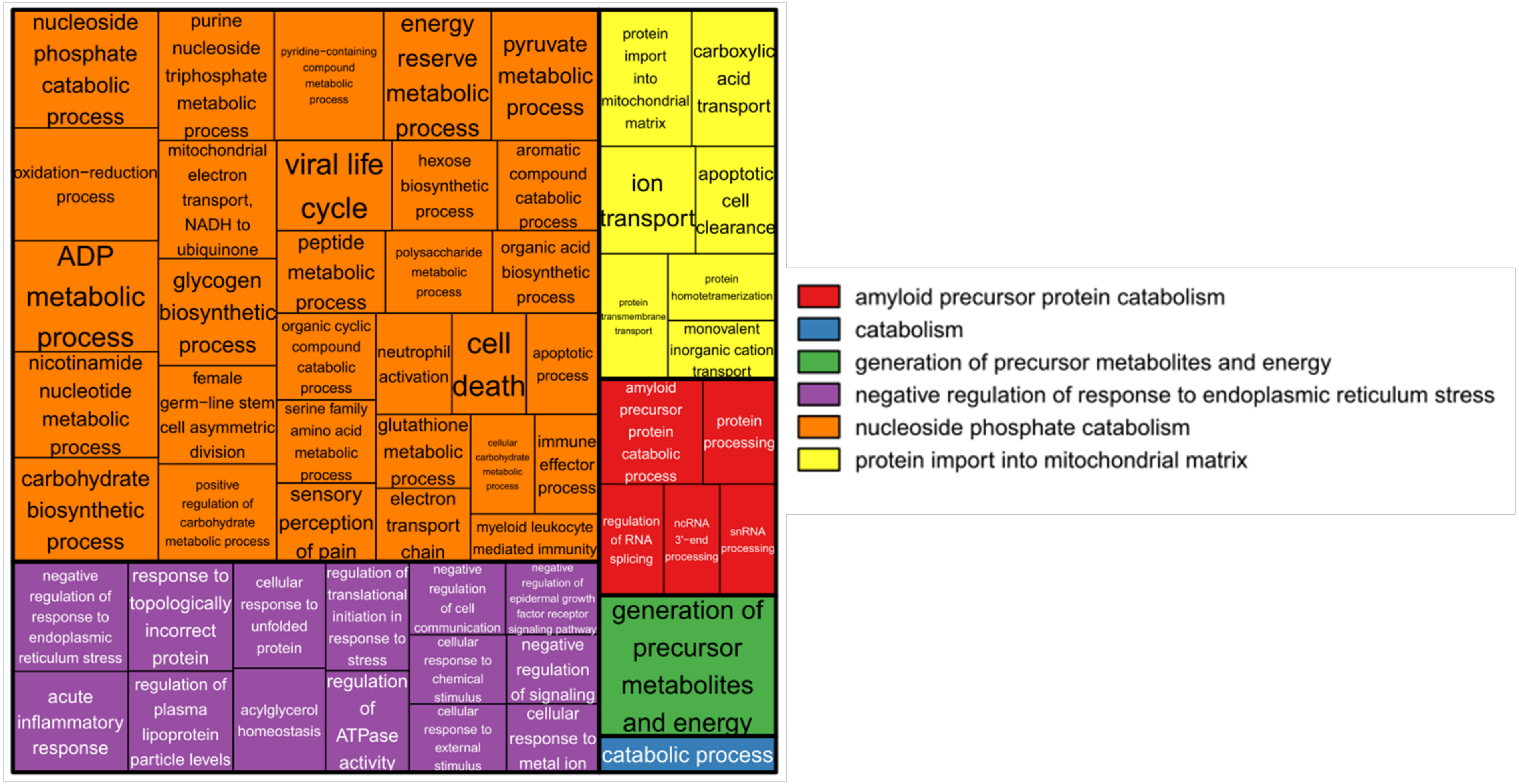


Figure S5: GO term enrichment for Biological Processes (BP). Enriched BP for GO terms ($p < 0.05$) associated with three or more significant differentially expressed genes, clustered using REVIGO. These rectangles are joined into different coloured 'superclusters' of loosely related terms. The area of the rectangles represents the p-value associated with that cluster's enrichment.
